# Design-driven optimization of low-cost reagent formulations for reproducible and high-yielding cell-free gene expression

**DOI:** 10.1101/2025.08.01.668204

**Authors:** Meagan L. Olsen, Caroline E. Copeland, Chad A. Sundberg, Rochelle Aw, Zachary M. Shaver, Govind Rao, James R. Swartz, Ashty S. Karim, Michael C. Jewett

## Abstract

Access to recombinant proteins is vital in basic science and biotechnology research. Cell-free gene expression systems provide one approach to address this need, but widespread utilization remains limited by the cost, complexity, and inconsistency of current platforms. To address these limitations, we carry out a multi-dimensional definitive screening design to reduce the number of reagent components and remove costly secondary energy substrates. From more than 1,200 reagent formulations, we discover a simple and reproducible system based on 12 components. The optimized reagent formulation can produce 2.4 ± 0.3 g/L of protein product at the 15-μL scale (∼$55/g_protein_) and 3.7 ± 0.2 g/L (∼$36/g_protein_) at the 4-mL scale with oxygen supplementation. This provides an 84 to 99% reduction in cost over previous cell-free reagent formulations. We further show that the optimized reagent formulation can produce nucleoside triphosphates from nucleotides and ribose and that it is robust to failure across batches of cell lysates, users/locations, and in the synthesis of different proteins. Specifically, we demonstrate the production of fifteen therapeutically relevant products, including full-length aglycosylated monoclonal antibodies. We anticipate that our optimized reagent formulation will further democratize the use of cell-free systems for protein manufacturing and synthetic biology applications.

## Introduction

Cell-free gene expression (CFE) systems can produce recombinant proteins in cell lysates upon incubation with essential substrates (e.g., amino acids, DNA template, energy substrates).^1–3^ These systems have advanced applications in diagnostics^4–15^, pathway prototyping^16–21^, protein design^22–26^, metabolic engineering^27–31^, and education^32–36^. Cell-free systems provide numerous benefits for protein production that are complementary to cell-based production platforms. First, cell-free systems are modular due to their open reaction environment; new functionalities (e.g., post-translational modifications) can be added through drop-in reagents. In addition, new protein products can be made using the same biocatalyst (i.e., same cell lysate) by changing the encoded DNA added to the reaction; the lysate source strain does not need to be re-engineered. Second, cell-free systems are scalable from the nL to the 1,000-L scale^37^ with growth-independent batch-to-batch performance. Lastly, distributed manufacturing paradigms can be enabled by cell-free systems, which can be freeze-dried, distributed, stored, and then readily reactivated by just adding water^38–41^. State-of-the-art cell-free expression systems can now produce a wide variety of biologics, including full-length aglycosylated monoclonal antibodies and glycosylated therapeutics^22,26,38–40,42–52^.

Despite the growing use of cell-free systems, reagent costs remain expensive for widespread commercial adoption. Cell-free gene expression, for example, depends on small-molecule reagent components that include energy substrates (e.g., phosphoenolpyruvate); DNA, RNA, and protein building blocks (e.g., NTPs, amino acids); cofactors and coenzymes (e.g., CoA, NAD); salts (e.g., magnesium, polyamines); and other chemical components (e.g., buffers).

Many reagent formulations rely on expensive phosphorylated energy substrates (e.g., $2,000- 3,000/L_CFE_), such as phosphoenolpyruvate (PEP), to regenerate the ATP that provides energy for protein biosynthesis^53–55^. This results in reagent costs upwards of $4,000/L_CFE_ and contributes to over 75% of all material costs associated with cell-free expression (**Table 1; Table S1**). While one report suggests cell-free systems can produce upwards of 4 g/L of protein in batch operation^56^, most studies report < 2 g/L of protein^1^, which leaves costs an order of magnitude higher than the $10s-$100s/g_protein_ associated with cell-based production^57,58^. Manufacturer availability and variability of complex reagents like purified tRNA further complicate implementation.

**Table 1.** A comparison of cell-free reagent formulations. All numbers indicate concentration in mM, except for folinic acid, tRNA, and maltodextrin, which are given in mg/mL, and costs and yields, which have units listed.

|  | Previously Published Systems |  |  |  |  |  |  |  | This Work |  |
| --- | --- | --- | --- | --- | --- | --- | --- | --- | --- | --- |
|  | Jewett and Swartz: PANOx-SP (2004) | Jewett and Swartz (2004) | Calhoun and Swartz (2005) | Zawada, et al. (2011) | Cai, et al. (2015) | Garenne, et al. (2021) | Pandi, et al. (2022) | Warfel, et al. (2023) | Minimal | RF<sub>opt</sub> |
| CFE cost (\$<sub>reagent</sub>/g<sub>protein</sub>) | \$3,760 | \$3,177 | \$2,782 | \$335 | \$556 | \$4,547 | \$7,763 | \$350 | \$165 | \$55 |
| Reagent cost (\$/L<sub>CFE</sub>) | \$4,513 | \$1,954 | \$1,906 | \$304 | \$253 | \$4,574 | \$4,316 | \$253 | \$87 | \$133 |
| Yield determined in this work (g/L) | 1.11 ± 0.13 | 0.62 ± 0.02 | 0.72 ± 0.03 | 0.91 ± 0.08 | 0.45 ± 0.03 | 1.01 ± 0.10 | 0.56 ± 0.02 | 0.72 ± 0.05 | 0.53 ± 0.01 | 2.39 ± 0.28 |
| Reported yield (g/L) | 0.7 | 0.7 | 0.5 | 0.7 | 1.3 | 4.0 | 0.5 | 1.2 | - | - |
| Mg(Glu)<sub>2</sub> | 8 | 8 | 8 | 8 | 8 | 8 | 10 | 10 |  | 8 |
| NH<sub>4</sub>(Glu) | 10 | 10 | 10 | 10 |  |  |  | 10 |  |  |
| K(Glu) | 130 | 130 | 130 | 130 | 260 | 80 | 105 | 130 | 300 | 362 |
| Glucose |  |  | 30 |  |  |  |  |  |  | 10 |
| Amino acids | 2 | 2 | 2 | 2* | 2* | 1.5** | 0.87 | 2 | 3.25 | 5 |
| Potassium phosphate |  |  | 10 | 15 | 15 |  |  | 75 |  | 15 |
| Nicotinamide |  |  |  |  |  |  |  |  |  | 4 |
| Ribose |  |  |  |  |  | 30 |  |  |  | 50 |
| HEPES | 57 |  |  |  |  | 50 | 50 |  |  | 75 |
| Bis-Tris |  |  | 57 |  |  |  |  | 57 |  |  |
| Pyruvate |  | 33 |  | 35 |  |  |  |  |  |  |
| Putrescine | 1 | 1 | 1 | 1 |  |  |  | 1 |  |  |
| Spermidine | 1.5 | 1.5 | 1.5 | 1.5 | 1.5 | 1 | 1 | 1.5 |  |  |
| Dithiothreitol |  |  |  |  |  | 1 | 2.5 |  |  |  |
| Folinic acid | 0.03 | 0.034 | 0.034 |  |  | 0.032 | 0.068 | 0.034 |  |  |
| tRNA | 0.17 | 0.17 | 0.17 |  |  | 0.2 | 0.04 |  |  |  |
| CoA | 0.27 | 0.26 | 0.26 |  |  | 0.26 | 0.13 |  |  |  |
| NAD | 0.4 | 0.33 | 0.33 |  |  | 0.33 | 0.175 | 0.4 |  |  |
| cAMP |  |  |  |  |  | 0.75 | 0.65 |  |  |  |
| PEP | 30 |  |  |  |  |  |  |  |  |  |
| 3-PGA |  |  |  |  |  | 30 | 28 |  |  |  |
| Oxalic acid | 4 | 4 |  | 4 | 4 |  |  | 4 |  |  |
| GSSG |  |  |  | 4 | 2 |  |  |  |  |  |
| GSH |  |  |  | 1 |  |  |  |  |  |  |
| Maltodextrin |  |  |  |  |  | 21.6 |  | 60 |  |  |
| PEG-8000 |  |  |  |  |  |  | 4 |  |  |  |
| ATP | 1.2 | 1.2 | 1.2 |  |  | 1.5 | 1.56 |  |  |  |
| CTP | 0.85 | 0.85 | 0.85 |  |  | 0.9 | 1.56 |  |  |  |
| GTP | 0.85 | 0.85 | 0.85 |  |  | 1.5 | 1.56 |  |  |  |
| UTP | 0.85 | 0.85 | 0.85 |  |  | 0.9 | 1.56 |  |  |  |
| AMP |  |  |  | 1.2 | 1.2 |  |  | 1.2 | 1.2 | 3 |
| CMP |  |  |  | 0.86 | 0.86 |  |  | 0.86 | 0.86 | 2.15 |
| GMP |  |  |  | 0.86 | 0.86 |  |  | 0.86 | 0.86 | 2.15 |
| UMP |  |  |  | 0.86 | 0.86 |  |  | 0.86 | 0.86 | 2.15 |

\*1 mM tyrosine. \*\*1.25 mM leucine.

To address high costs of cell-free gene expression, numerous groups have developed different reagent formulations that leverage less expensive non-phosphorylated energy secondary substrates for reagent costs of < $500/L_CFE_ (**Table 1**)^39,43,53,59–62^. However, these reagent systems have not been widely adopted. As such, reducing costs while improving protein yields remains essential for commercial use of cell-free protein manufacturing strategies.

In this work, we aimed to develop a cell-free expression reagent formulation that enables low-cost, consistent, and high-yield protein production. A key feature of our approach was the focus on reducing the number of components used while simultaneously designing for yield and cost metrics. The goal was to reach less than $100 in reagents per gram of protein to more closely match cell-based production costs^57,58^. Inspired by previous multi-dimensional optimization efforts^63,64^, we carried out design-driven optimization to screen 58 cell-free reagent components and their concentrations in 1,231 different reaction combinations. Our most productive formulation was able to synthesize 2.4 ± 0.3 g/L superfolder green fluorescent protein (sfGFP) at a reagent cost of $133/L_CFE_ ($55/g_protein_) in a standard batch reaction. Improving oxygen flux further increased yields to 3.7 ± 0.2 g/L sfGFP ($36/g_protein_). This optimized formulation lowers cell-free expression cost by an average of 95% over state-of-the-art phosphorylated energy substrate formulations. Lastly, we demonstrate modularity for protein manufacturing by using this reagent formulation to produce fifteen recombinant protein therapeutics, including full-length aglycosylated monoclonal antibodies and other disulfide bonded products. We anticipate that low-cost, consistent, and high-yield cell-free production will expand existing biomanufacturing efforts, helping to meet the need for recombinant proteins.

## Results

### Comparing state-of-the-art cell-free reagent formulations

We first set out to create a baseline comparison of previously reported reagent formulations by experimentally examining their protein production capacity in our laboratory (**Table 1**). By doing this, we can fairly compare the value of reagent differences in these formulations. Specifically, we tested eight previously reported reagent formulations for *E. coli* crude lysate platforms^39,43,53–55,59,60^, focusing on low-cost variants that leverage non-phosphorylated energy substrates like glucose or glutamate. We used an *E. coli* BL21 Star (DE3) lysate to test each formulation, though some of the formulations were originally designed around other B- and K-strains of *E. coli*. We individually assembled each formulation, constructed 15-μL cell-free gene expression reactions containing pJL1-sfGFP plasmid DNA, and incubated reactions at 30 °C for 20 h. We found that the PANOx-SP formulation (our phosphorylated energy substrate base case)^53^ produced the highest sfGFP yield and used this formulation as a standard for the remainder of the work. Although all non-phosphorylated energy substrate systems produced less total protein than the phosphorylated substrate formulations, the reagent cost per gram of protein ranged from 22-92% less than that of the PANOx-SP system (**Table 1**; **Fig. 1a**).

**Figure 1.**
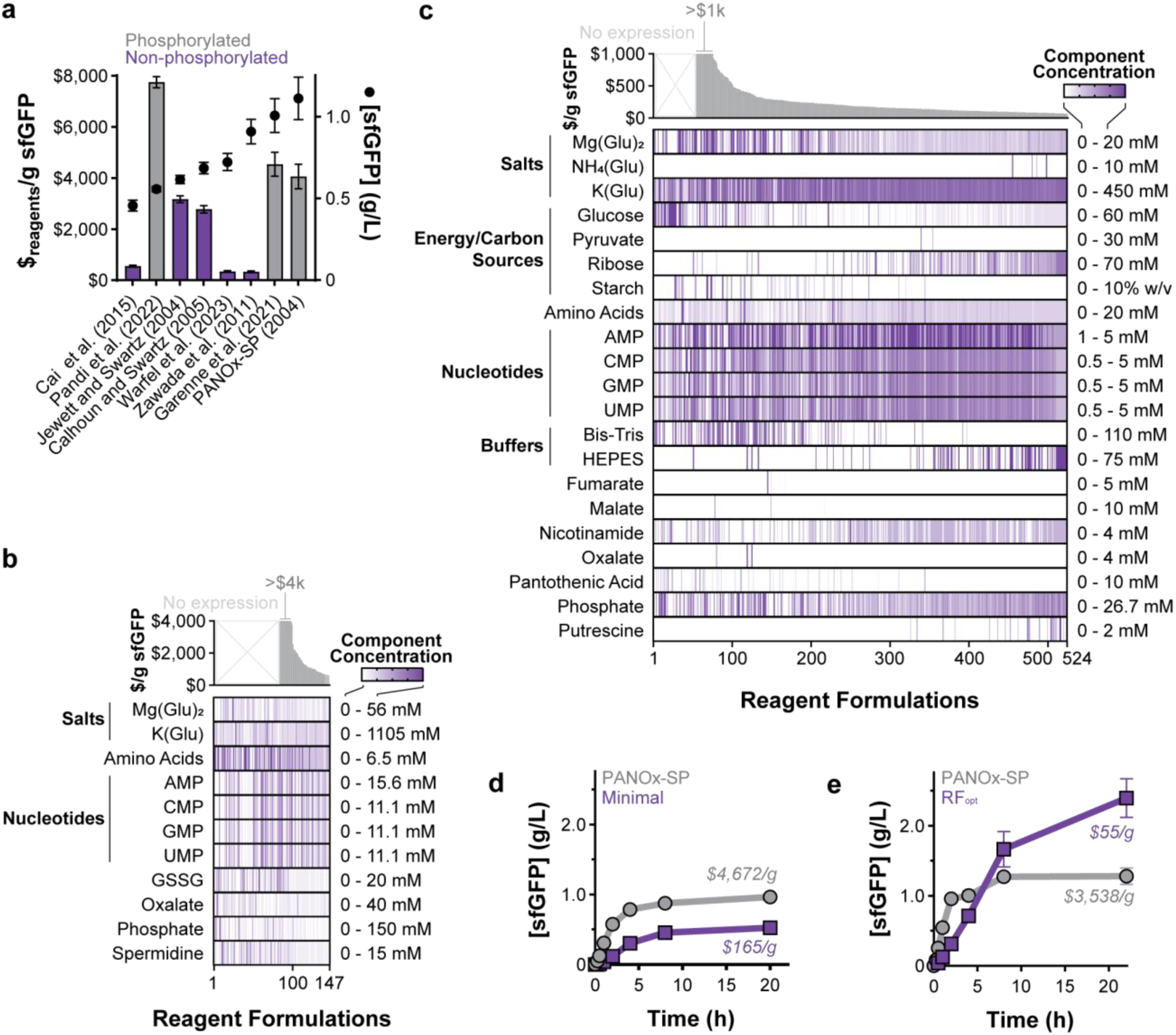
Development of a low-cost and high-yielding reagent formulation. (a) sfGFP yields (dots) and cost of reagents (bars) per gram sfGFP produced were determined for 8 cell-free reagent formulations. Costs are based on only raw materials purchased at the laboratory scale, as calculated in **Table 1** and **Table S2**. Error bars show standard deviation (g/L) and propagated error ($/g) for *n* = 3 sfGFP expression replicates. (b) Reagent concentrations sorted by their corresponding $/g_protein_ for the initial screen of low-cost cell-free reagent components. Color gradients are used to represent component concentrations for each tested reagent formulation. (c) Reagent concentrations sorted by their corresponding $/g_protein_ for the multi-component optimization campaign, building off the reagent formulation in (b). NMP concentrations were capped at 5 mM to prevent formation of insoluble precipitates. (d) Cell-free expression levels over 22 h using either the PANOx-SP or minimal reagent formulations. Reactions were performed in a 384-well plate. (e) Cell-free expression levels over 22 h using either the PANOx-SP or optimized reagent formulation (RF_opt_). Reactions were performed in 2-mL flat-bottomed Axygen tubes. Error bars for panels (d) and (e) represent the standard deviation of n = 3 replicates. All reactions were run using BL21 Star (DE3) lysate at 30 °C for 20-22 h.

#### Optimizing a cell-free reagent formulation

Building on these established reagent formulations, we set out to identify a low-cost, consistent, and high-yielding cell-free reagent formulation (i.e., low $_reagent_/g_protein_). We began with a panel of 11 reagent components common amongst the established reagent formulations. These components included magnesium glutamate, potassium glutamate, the 20 standard amino acids (counted as a single component), phosphate, oxalate, spermidine, and oxidized glutathione (**Fig. 1b**). We used a combination of definitive screening designs, targeted reagent range adjustments, and stepwise models built from the compiled data using JMP software to screen these components. This initial analysis of 147 reactions identified an active reagent formulation comprised only of potassium glutamate, nucleotide monophosphates, and amino acids. A three-factor optimization yielded a formulation that produced 0.5 ± 0.1 g/L sfGFP (**Fig. 1d; Fig. S1**). This minimal system ($165/g_protein_) is the smallest set of reagent components reported for any *in vitro,* combined transcription and translation system.

Next, we explored individual supplementation of other reagents canonically used in cell-free gene expression to our minimal system to see if we could improve total protein yields while keeping the number of reagents low. We constrained our search to lower cost reagents (i.e., less than $50/g_reagent_) to balance the need for low-cost formulations with potential increased protein yields. Many of the more expensive reagents, including tRNA, coenzyme A, and NAD^+^, have previously been shown to be unnecessary for cell-free gene expression when cell lysates are not dialyzed^39,43^, so we did not include them here. We found that individual additions of buffer (e.g., HEPES, Bis-Tris), magnesium glutamate, alternative energy substrates (e.g., glucose, pyruvate, maltodextrin, ribose), oxalate, spermidine, phosphate, and TCA cycle intermediates (e.g., fumarate, malate) did not improve protein yield (**Fig. S2**). Moreover, individual replacements of nucleoside monophosphates—the most expensive reagent group ($62/g_protein_) in the minimal formulation—with their corresponding nucleosides or nucleobases generally decreased protein yields (**Fig. S3**). Insoluble precipitates were also observed when using many of these alternatives, suggesting further investigation into precipitate formation and prevention might be necessary to evaluate these modifications. Surprisingly, we found that GMP could be replaced with a combination of guanine and ribose without sacrificing protein yield. These results and known synergistic benefits between reagents (e.g., glucose and phosphate^60^) suggested that any significant improvement to protein yield would require simultaneous supplementation of multiple reagents.

Inspired by previous multi-dimensional reagent optimization efforts^63,64^, we next used a combination of definitive screening designs and targeted reagent manipulations to explore simultaneous addition of multiple reagents to the minimal formulation (**Fig. 1c**). We included all individually tested reagents as well as additional salts (i.e., NH ^+^), energy alternatives (i.e., starch), nicotinamide, pantothenic acid, and putrescine—21 components in total. We leveraged stepwise models based on the collected data to identify and test putative optimum conditions. Best-performing formulations were then used as the baseline for further rounds of definitive screening designs as well as targeted optimization based on learned design rules, such as yield improvement through addition of ribose and HEPES. Through this iterative process, we explored the design space of 2^21^ (i.e., 2,097,152) reagent combinations with only 524 formulations, increased protein yield to over 2 g/L sfGFP, and drove down reagent costs 15-fold (**Fig. S4**). The best performing minimal reagent formulation produces 2.4 ± 0.3 g/L sfGFP within 20 h (**Fig. 1e**). This is an 87% increase in protein yield over the previously best-performing PANOx-SP system. The optimized reagent formulation (RF_opt_) costs $133/L_CFE_ ($55/g_protein_), 99% less than the PANOx-SP system and 84% less than the previous lowest cost system—a 2 orders of magnitude cost reduction (**Table 1**).

#### Characterizing the optimized reagent formulation

To better understand the behavior of our optimized minimal reagent formulation, we compared cell-free metabolism over the course of a 20-h cell-free expression reaction using the PANOx- SP formulation and our optimized reagent formulation (RF_opt_). We first observed ATP and phosphate profiles as these have been shown to be key measures of cell-free gene expression performance^31,65,66^. In line with previous works using the PANOx-SP formulation^65^, a surge of ATP is produced within 30 minutes with a corresponding rapid and sustained increase in phosphate as a result of the phosphorylated energy substrate PEP being consumed (**Fig. 2a-b**). ATP levels then drop to nearly zero by the end of the reaction. RF_opt_ generates comparable levels of ATP within 30 minutes, despite starting with no exogenous ATP, and sustains ATP levels at ∼250 μM (well above the ATP affinity threshold for cell-free gene expression^66^) for the duration of the reaction (**Fig. 2a**). Phosphate levels hover around 10-15 mM over the reaction lifetime, three times lower than the PANOx-SP formulation, because free phosphate is not liberated from PEP as in the PANOx-SP system (**Fig. 2b**). Furthermore, we measured key central carbon metabolites (pyruvate, lactate, acetate, citrate, succinate, fumarate, and malate) that often serve as carbon and energy intermediates and sinks in metabolism for both formulations^60,67,68^ (**Fig. 3c; Fig. S5**). Time-course measurements show that RF_opt_ sustains the tricarboxylic acid cycle metabolites at much lower concentrations during the reaction compared to the PANOx-SP formulation. These results suggest that RF_opt_ balances cell-free metabolism for a longer period, allowing for more sustained rates of protein synthesis.

**Figure 2.**
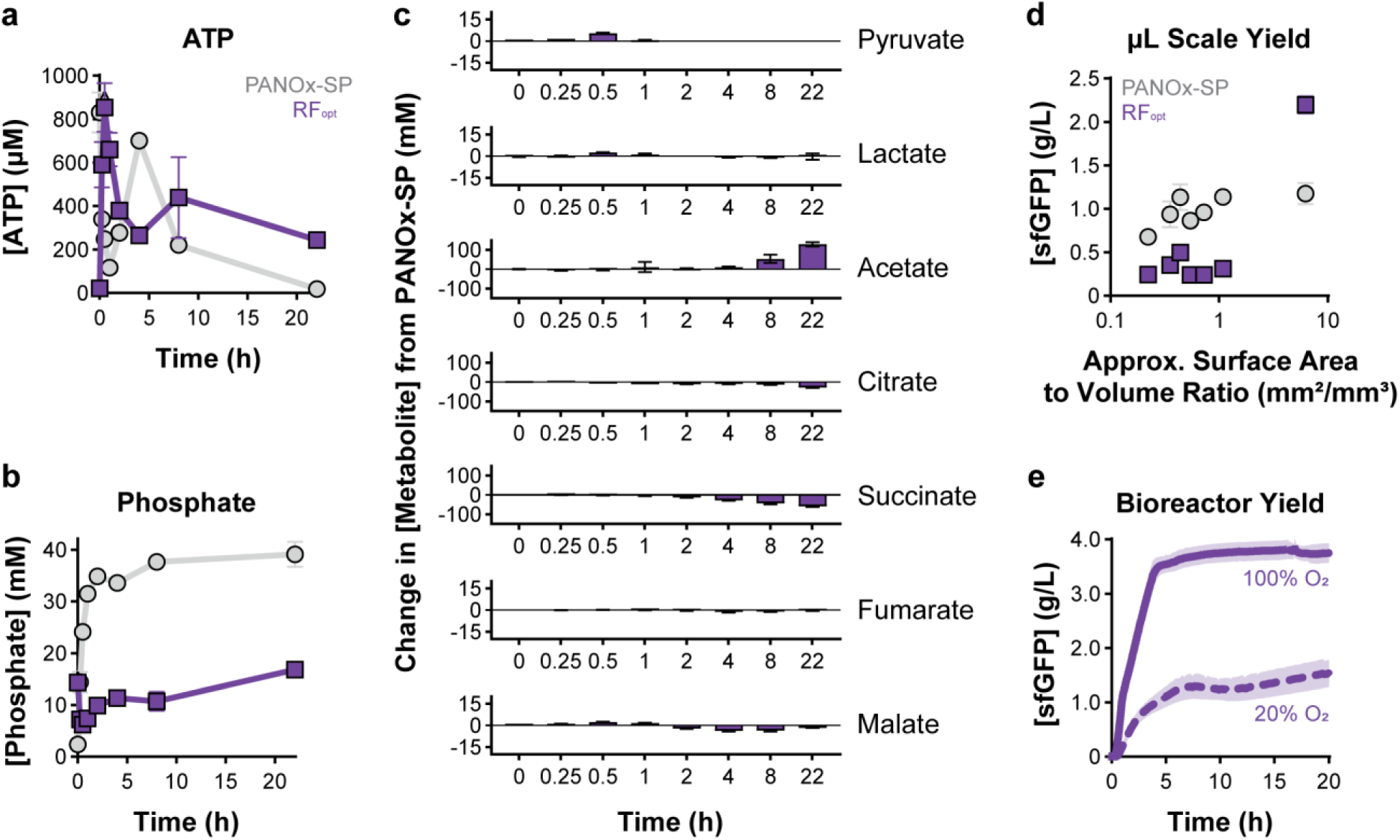
Adjusting reagent formulations alters the cell-free metabolic environment and oxygen dependency. (a) ATP and (b) phosphate concentrations in the cell-free reaction environment for the PANOx-SP and RF_opt_ formulations. Error bars represent the standard deviation of *n* = 3 replicates. (c) Change in metabolite concentration between PANOx-SP and RF_opt_. Error bars represent the propagated error from the standard deviation of *n* = 3 replicates. (d) Comparison of sfGFP yield and approximate surface area to volume ratio (mm^2^/mm^3^), as determined by manufacturer tube/plate schematics and caliper measurements. Error bars represent the standard deviation of *n* = 3 replicates. (e) sfGFP production in a 4-mL bioreactor supplemented with either a 20% or 100% O_2_ feed. The shaded area represents standard deviation for *n* = 6 bioreactors with a 20% O_2_ feed and *n* = 3 bioreactors with a 100% O_2_ feed. Additional replicates were run for the 20% O_2_ feed due to the increased variability. All reactions were run using *E. coli* BL21 Star (DE3) lysate at 30 °C for 20-22 h.

**Figure 3.**
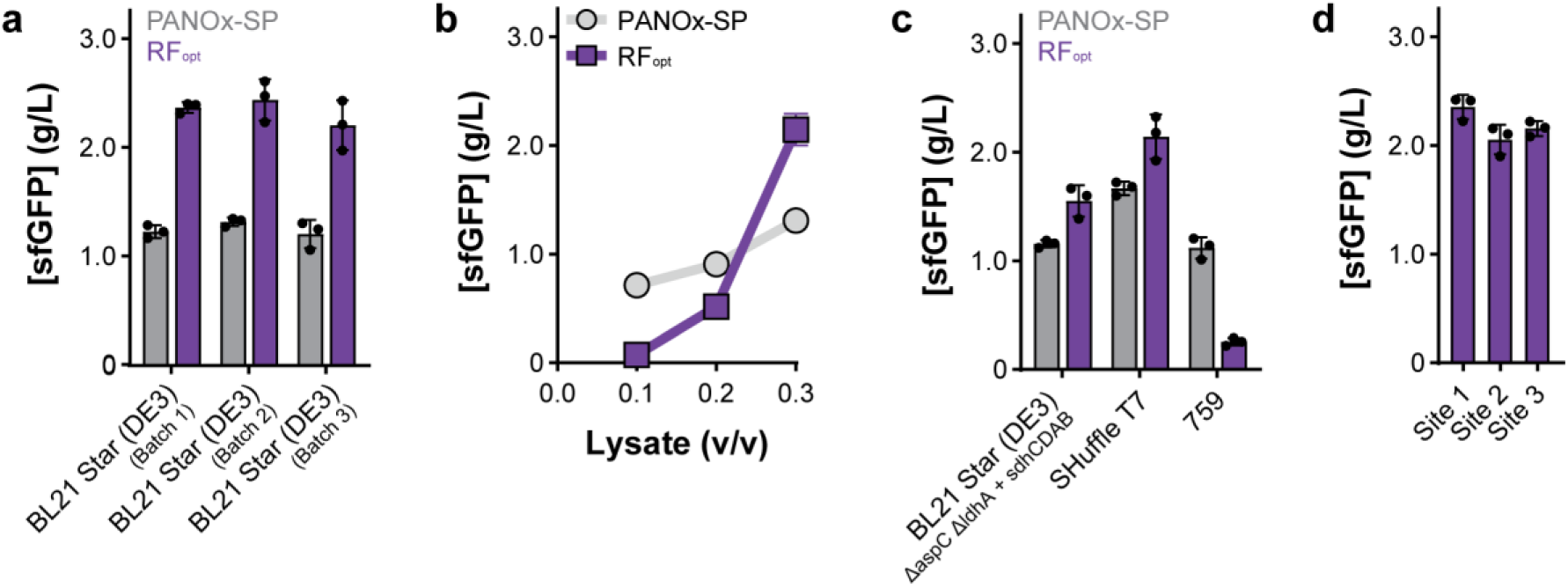
The optimized reagent formulation (RF_opt_) remains robust across batches of cell lysates and users/locations. (a) Productivity of the PANOx-SP and RF_opt_ formulations across three independent *E. coli* BL21 Star (DE3) lysate batches. (b) Impact of lysate concentration on sfGFP expression level. New optimum magnesium and phosphate concentrations were determined for each lysate concentration. (c) sfGFP expression in three additional *E. coli* lysates with optimized magnesium and phosphate concentrations. (d) Protein production using RF_opt_ at three different laboratory sites, each using reagents and lysate prepared independently. In all panels, error bars show standard deviation for *n* = 3 replicates. All reactions were run at 30 °C for 20-22 h.

Next, we investigated the performance of RF_opt_ across different reaction geometries, known to be important for oxygen transfer^69,70^. To do this we tested several reaction vessels (i.e., 2-mL tubes, 0.2-mL tubes, 384-well plates) and volumes (i.e., 15-100 μL) to vary the surface area-to-volume ratio between 0.1 and 10 (**Fig. 2d; Fig. S6**). We found that cell-free expression using the optimized formulation had greater sensitivity to this ratio, suggesting oxygen levels may limit reaction yields. We leveraged a 4-mL membrane-based bioreactor^71^ with a tunable oxygen feed to further examine this behavior (**Fig. S7**). When provided with a 20% O_2_ feed to mimic ambient air, the bioreactor-housed cell-free gene expression reaction produced less than 2 g/L sfGFP (**Fig. 2e**). Shifting to a 100% O_2_ feed increased the dissolved oxygen (dO_2_) content in the reaction (**Fig. S7**; **Fig. 2e**) and improved sfGFP production by 45%, achieving 3.7 ± 0.2 g/L sfGFP ($36/g_protein_) at the 4-mL scale. Protein synthesis is also more rapid, producing 0.9 ± 0.1 g/L/h for the first four hours in the bioreactor. In contrast, the 20% O_2_ feed produced roughly 0.3 ± 0.04 g/L/h and the 15-μL tube-based reaction produced 0.2 ± 0.01 g/L/h. These results suggest that control of metabolism and oxygen transfer will be key to extending reaction rate, longevity, and yield.

#### Evaluating robustness of an optimized cell-free reagent formulation

Optimal reagent formulations must be robust to failure across batches of cell lysates, users/locations, and in the synthesis of different proteins for widespread use. To test robustness, we performed a series of experiments measuring sensitivity to these important parameters. We evaluated our optimized reagent formulation (RF_opt_) using various cell lysate batches, volumes, and source strains. First, we grew three independent cultures of *E. coli* BL21 Star (DE3) and made subsequent lysates as described in **Methods**. Magnesium and phosphate concentrations were optimized for each new lysate (**Fig. S8**). Running cell-free expression reactions using these lysates and RF_opt_ produces consistent protein yields across these three lysate batches (**Fig. 3a**). Next, we found that protein yields are more sensitive to the amount of lysate added than using the PANOx-SP formulation (**Fig. 3b**), reiterating the fine metabolic balance we observed (**Fig. 2**). We then assessed RF_opt_ for cell-free expression with lysates made from other strains of *E. coli*. We specifically evaluated three other strains: (i) a genomically modified variant of BL21 Star (DE3) to test sensitivity to typical metabolic modifications; (ii) SHuffle T7, another B-strain commonly used with enhanced capacity to correctly fold proteins with disulfide bonds; and (iii) a K-strain *E. coli*, 759, engineered for expanded genetic code applications^23,72^. Following lysate preparation, we ran cell-free expression reactions with RF_opt_ (**Fig. 3c; Fig. S8**). We found that RF_opt_ significantly improves protein yields in B-strain-derived lysates compared to the PANOx-SP formulation, whereas RF_opt_ decreases protein yields in a K-strain derived lysate, requiring additional optimization as seen before^23,72^. We also found that RF_opt_ is robust to the addition of common salts, osmolytes, and protein buffers (**Fig. S9**), enabling use of the formulation across a variety of workflows.

In addition to a reagent formulation needing to be robust across batches of cell lysates, cross-laboratory reproducibility is essential for wider adoption of cell-free gene expression. While multi-site implementation has proven challenging^73,74^, we wanted to test RF_opt_ across different locations. To do this, we provided the RF_opt_ formulation to two other academic institutions, with a different researcher preparing reagents and setting up reactions at each site. We found that RF_opt_ produced protein yields within ∼9% between sites (**Fig. 3d**), demonstrating that this formulation is robust across users/locations.

Lastly, we wanted to ensure that RF_opt_ could be used effectively to synthesize a variety of commercially relevant protein products. We first chose to produce FDA-approved carrier proteins (protein D and CRM197) used in conjugate vaccines as well as the trastuzumab monoclonal antibody (both Fc domain and full-length antibody) used to treat HER2-positive breast and stomach cancers as model proteins of interest. While RF_opt_ produced some soluble product, it was less productive than the PANOx-SP formulation. However, neither condition produced the trastuzumab Fc dimer because oxidizing conditions are required to form disulfide bonds (**Fig. 4a; Fig. S10**). To form disulfide bonds, cell-free reactions use pretreatment with iodoacetamide (IAM) to deactivate reductases, addition of glutathione (GSSG/GSH) to create an oxidizing reaction environment, and supplementation of disulfide bond isomerase (e.g., DsbC) to aid in disulfide bond shuffling^45,75,76^. Surprisingly, the addition of these components to RF_opt_ significantly reduced sfGFP production by 75-97% (**Fig. 4b**). To mitigate indiscriminate IAM activity,^77^ we knocked out a glutathione reductase gene (Δ*gor*) in the lysate source strain to help stabilize the oxidizing lysate environment (**Supplemental Data File 1**), which recovered protein yields with 20-fold less IAM (**Fig. 4b; Fig. S11**). However, adding glutathione at several oxidized:reduced ratios still decreased protein yields by up to 62% (**Fig. 4b; Fig. S12**), without negatively impacting cell-free energy generation (**Fig. S13**). Diving deeper, we found that glutathione addition correlated with a decrease in reaction pH for both the RF_opt_ and the PANOx- SP formulations (**Fig. 4c**). While this is not problematic for the PANOx-SP formulation, which remains within the cytoplasmic pH of 7.4-7.8 ^78^, RF_opt_ is more acidic and the addition of glutathione rapidly drops the reaction pH below pH 7.0 (**Fig. 4c**). By increasing the HEPES buffer pH to 7.5, we avoided the pH drop and recovered protein yield (**Fig. 4c**; **Fig. S14**).

**Figure 4.**
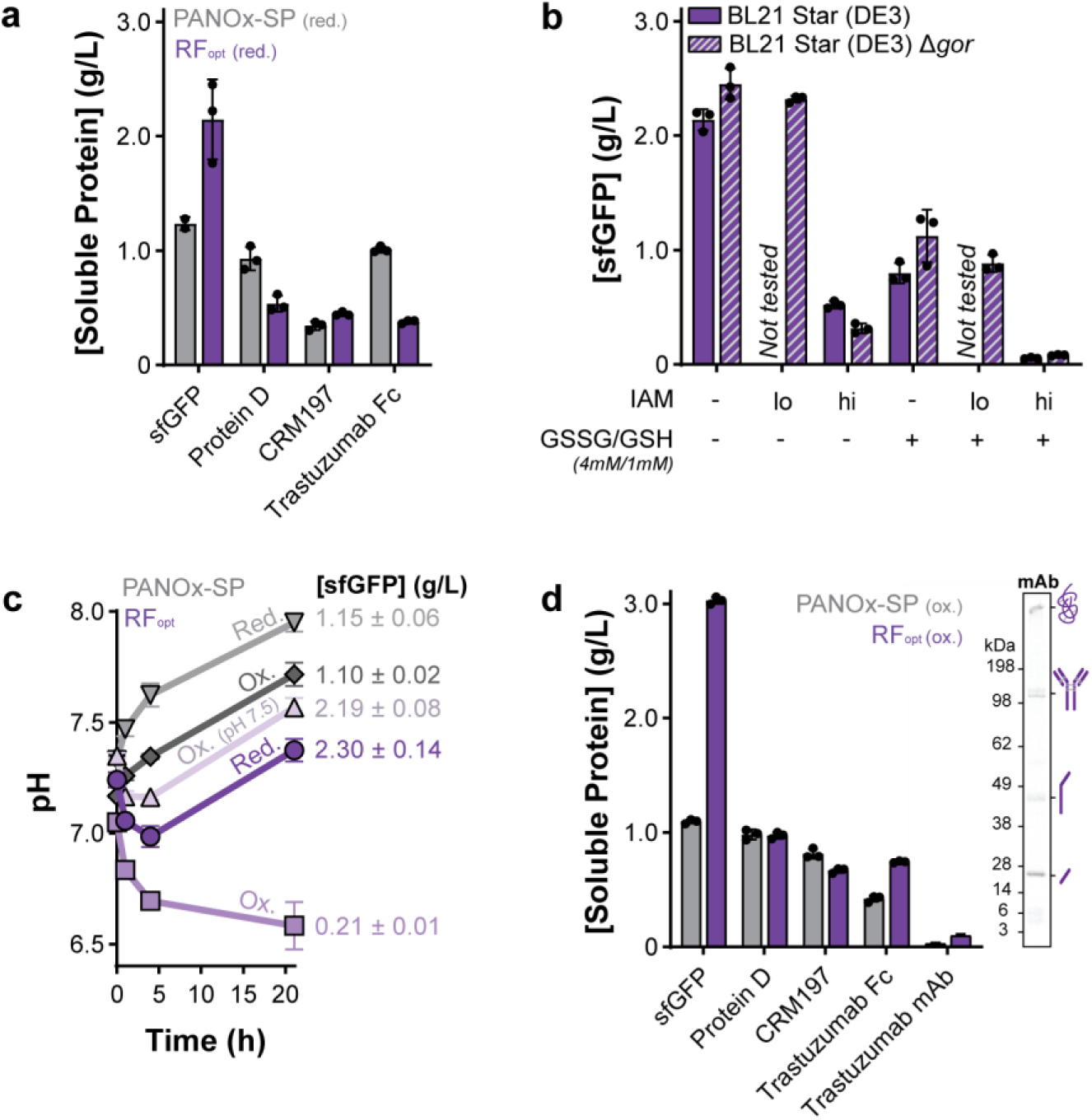
An oxidized variation of the optimized reagent formulation (RF_opt_) produces full-length aglycosylated antibody. (a) Soluble protein production in the unmodified PANOx-SP and RF_opt_ formulations using a BL21 Star (DE3) lysate. (b) Impact of oxidized/reduced glutathione (GSSG/GSH), iodoacetamide (IAM; lo = 25 μM, hi = 500 μM), and Δ*gor* genome modification on sfGFP yields when using RF_opt_. (c) pH traces and corresponding yields for a 21-h reaction when using the PANOx-SP or RF_opt_ formulations either in their unmodified reducing condition (red.) or with the addition of IAM, GSSG, and GSH (ox.). Unless otherwise noted, the HEPES buffer included in each reaction was initially at pH 7.2. (d) Soluble protein production in the oxidized versions of the PANOx-SP and RF_opt_ formulations. The insert shows the autoradiogram for trastuzumab full-length antibody (mAb) formation in the RF_opt_ system. A BL21 Star (DE3) Δ*gor* lysate was used in panels (c)-(d). All reactions were run at 30 °C for 20-22 h. Error bars represent the standard deviation of *n* = 3 replicates.

The oxidized version of RF_opt_ improved production of protein D, CRM197, and the trastuzumab Fc and full-length antibody constructs (**Fig. 4d**), increasing the fraction of assembled Fc domains and full-length antibodies by 44% and 90%, respectively, compared to an oxidized PANOx-SP reaction (**Fig. S15**). We then examined the preliminary IgG1 light chain expression to further improve full-length trastuzumab folding. By lowering the reaction temperature^70,79^ from 30 °C to 22 °C and delaying the addition of DNA encoding the heavy chain^44^ for 3-h, we produced ∼150 μg/mL full-length trastuzumab using RF_opt_ (**Fig. S16**).

#### Diverse therapeutic protein production in optimized reagent formulation

We next expressed fifteen medically relevant recombinant proteins and compared product yields to the most productive (PANOx-SP) and lowest $/g sfGFP cost reagent formulations^59^ (**Fig. 5a**). The selected proteins span 16-147 kDa, 0-16 disulfide bonds, and a variety of clinical applications, with six previously expressed in *E. coli*-based cell-free expression systems^24,38,40,43,50,76^. All protein yields were determined via ^14^C-leucine incorporation in an oxidized cell-free reaction environment with added bacterial DsbC and verified with autoradiography (**Figs. S17-S20**). We observed expression of all 15 therapeutic proteins with our optimized reagent formulation (RF_opt_) and obtained >100 μg/mL soluble full-length protein for 12 proteins with at least one formulation. RF_opt_ yielded the same or more soluble protein than the previous most productive (PANOx-SP) formulation for 13 of 15 proteins and had lower $/g_protein_ costs for 11 of 15 proteins than the previous lowest $/g_protein_ cost formulation^59^ (**Fig. S21**). However, the level of improvement varied across proteins. For instance, caplacizumab was expressed with significantly higher yields using RF_opt_, while streptokinase and myoglobin showed reduced yields compared to the PANOx-SP formulation (**Fig. 5a**). In addition, while the ratio of antibody heavy chain and light chain varied across reagent formulation, the non-phosphorylated energy systems reduced insoluble aggregation of antibody constructs (**Fig. S22**).

**Figure 5.**
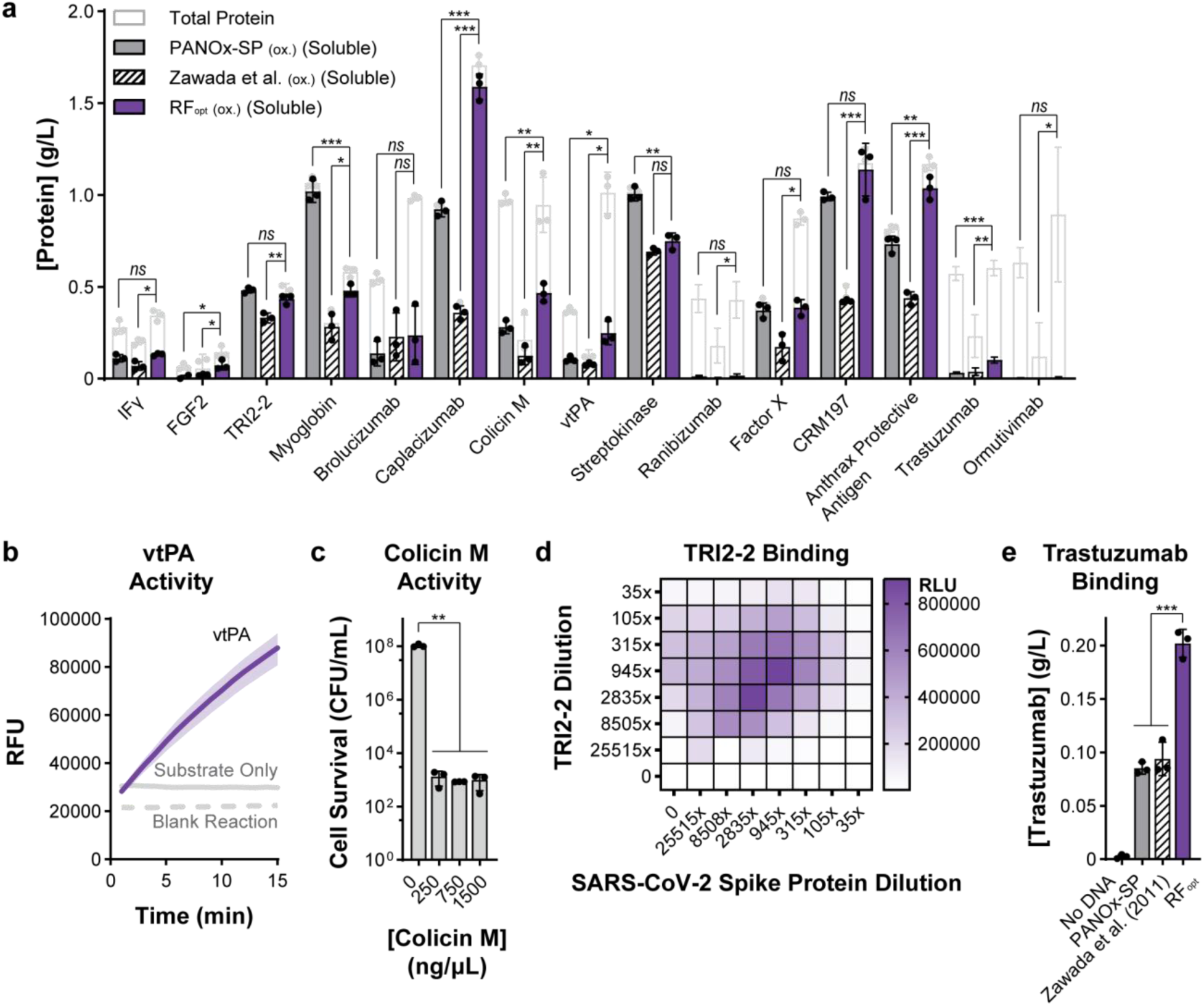
The optimized reagent system (RF_opt_) expresses 15 diverse recombinant protein therapeutics. (a) Expression of 15 recombinant protein therapeutics using three different reagent formulations, as quantified by ^14^C-leucine incorporation. All reactions were run in an oxidizing cell-free environment with BL21 Star (DE3) Δ*gor* lysate and included 5 μM of purified bacterial DsbC. Reactions were incubated at 30 °C for 20 h. Both total and soluble protein yields are shown for *n* = 3 replicates and error bars represent standard deviation. Statistical significance for soluble yields was calculated by unpaired two-tailed *t*-tests (adjusted *p* value < 0.001 denoted by ***, 0.001 to 0.01 by **, 0.01 to 0.05 by *, and > 0.05 by ns). (b) Enzymatic activity of vtPA produced using RF_opt_. The shaded area represents standard deviation for *n* = 3 replicates. The blank reaction contained all cell-free expression components except a DNA template. (c) Viability of CLM24 indicator cells treated with colicin M produced using RF_opt_. Error bars indicate standard deviation from *n* = 3 independent cell cultures. Statistical significance was calculated by an unpaired two-tailed *t*-test (adjusted *p* value < 0.001 denoted by ***, 0.001 to 0.01 by **, 0.01 to 0.05 by *, and >0.05 by ns). (d) AlphaLISA binding pattern generated by interaction of the cell-free produced TRI2- 2 minibinder and the SARS-CoV-2 spike protein. This experiment was run in duplicate, and all results can be found in Fig. S23. “0” indicates AlphaLISA reactions that do not contain any of the respective reaction component. (e) Full-length trastuzumab detected using an anti-idiotypic sandwich ELISA assay. Error bars indicate standard deviation from *n* = 3 replicates. Cell-free reactions used to generate protein for panels (b)-(e) were run in an oxidizing cell-free environment at 30 °C for 20 h using BL21 Star (DE3) Δ*gor* lysate supplemented with 5 μM purified bacterial DsbC. For trastuzumab production, heavy chain pDNA was added after 1.5 h of incubation. Statistical significance was calculated by an unpaired two-tailed *t*-test (adjusted *p* value < 0.001 denoted by ***, 0.001 to 0.01 by **, 0.01 to 0.05 by *, and >0.05 by ns).

Finally, we wanted to show that proteins expressed with RF_opt_ were functional. We selected a subset of our expressed proteins with a variety of functions—vtPA, colicin M, TRI2-2, and trastuzumab—to test for protein activity. vtPA is an enzymatically active truncated form of tissue plasminogen activator, which cleaves an arginine-valine peptide bond in plasminogen to activate the protease and dissolve blood clots^76,80^. We found that cell-free-expressed vtPA is functional and able to rapidly cleave the fluorescent 7-amino-4-methylcoumarin (AMC) from a short Ile-Pro-Arg peptide sequence^81^ (**Fig. 5b**). Next, we tested antimicrobial activity of colicin M^82^ by incubating cell-free-expressed colicin M with *E. coli* CLM24 indicator cells. The presence of colicin M decreased *E. coli* colony formation by a factor of 10^5^ compared to a no-DNA cell-free reaction matrix (**Fig. 5c**). Lastly, we tested TRI2-2 and trastuzumab function through binding assays. Using AlphaLISA—a protein-protein interaction assay that produces luminescence based on the proximity of donor and acceptor beads decorated with the proteins of interest^83^—we found cell-free-expressed TRI2-2, a computationally designed miniprotein inhibitor of SARS-CoV-2^24^, indeed binds to the SARS-CoV-2 spike protein to produce AlphaLISA signal 2.4-fold over background (**Fig. 5d, Fig. S23**). With an anti-idiotypic sandwich ELISA assay—trastuzumab antibody bound to a capture antibody designed to interact with the trastuzumab antigen binding site^84^—cell-free-expressed trastuzumab effectively bound to the capture antibody (**Fig. 5e**). These results demonstrate that RF_opt_ can robustly express active proteins with a variety of functions.

## Discussion

Cost and productivity are pivotal determinants for widespread use of cell-free systems for protein production. Here, we explored 1,231 reaction mixtures to establish a low-cost, high-yielding cell-free reagent formulation capable of producing >2 g/L protein product for $133/L_CFE_ in reagents. Notably, we identified a minimal reagent formulation containing only six components (with all 20 amino acids counted as one component), which comprises the fewest components of any reported formulation. We also developed an optimized minimal reagent formulation with 12 components. This optimized formulation (RF_opt_) reduces reagent cost from 75% of total material costs to less than 4% (**Table S1**).

The optimized reagent formulation (RF_opt_) has several key features. First, it has the highest reported protein yield for a reagent mixture leveraging non-phosphorylated energy substrates and the lowest reported cost to date (∼$55/g_protein_). A notable discovery was the importance of dissolved oxygen concentration on system performance; high DO_2_ levels increased yields to >3.7 g/L in a bioreactor, approaching the highest reported yields in a cell-free system (4 g/L)^56^ but for a fraction of the cost. Second, the formulation is robust to failure across batches of cell lysates, users/locations, and in the synthesis of different proteins. Third, the platform can produce proteins with disulfide bonds when pH is controlled. These features increase the economic viability of using cell-free expression for protein design^22,24,26^, enzyme engineering^21,25^, and therapeutic production^38,40,42,43^.

This work highlights the modular nature of cell-free systems, which allows for on-demand production of a diverse product library. Importantly, by just changing the DNA input, the same system formulation can be used to make different protein products. We showcased the optimized reagent formulation by synthesizing fifteen therapeutic proteins spanning 16-147 kDa, 0-16 disulfide bonds, and a variety of clinical applications (**Fig. 5**). Nine of these proteins had never been expressed in cell-free systems, and 12 proteins were produced at > 100 μg/mL soluble full-length protein. With these yields, cell-free systems could feasibly be integrated into point-of-care or distributed biologics manufacturing paradigms to complement centralized facilities that rely on cellular protein expression. Coupled with the large body of work on freeze-dried systems^39,47,85^, cell-free systems provide the necessary flexibility and reduction in transportation and storage costs for distributed, on-demand manufacturing^86–89^.

While our work brings us a step closer to realizing the vast potential of cell-free systems in distributed manufacturing, continued advances in purification strategies^42,90–92^ are required. Further reductions in cost, especially involving DNA template preparation^70,93^ and nucleoside monophosphates, and increases in productivity and longevity are also necessary. Our work highlights one potential path. Specifically, we demonstrated with our minimal formulation that a combination of guanine and ribose can replace GMP without a decrease in system performance (**Fig. S3**). These findings indicate that the metabolic pathways necessary to build and sustain full reagent profiles from a limited number of low-cost substrates are active and can be intentionally designed. Combining these results with advances in self-sustaining translation machinery^94,95^, DNA templates^96^, and amino acids^97^ may provide a path to a lower-cost and longer-lasting cell-free platform capable of building all components it needs to function.

In sum, the low-cost, high-yielding cell-free platform developed in this work offers a robust and versatile approach to rapidly produce recombinant proteins. By doubling yields of conventional cell-free technologies, we anticipate that the optimized reagent formulation will be adopted across numerous application spaces, from supporting protein design efforts to cell-free biomanufacturing at industrial scales for biologics production.

## Methods

### Cell Extract Preparation

Crude extracts were made from multiple *Escherichia coli* strains, listed in **Table S3**, based on past protocols^53,98,99^. In brief, cells were grown overnight at 37 °C in LB media (10 g/L tryptone, 5 g/L yeast extract, 5 g/L NaCl) to prepare a starter culture. The following day, 1 L cultures of sterilized 2xYTPG media (16 g/L tryptone, 10 g/L yeast extract, 5 g/L NaCl, 7 g/L K_2_HPO_4_, 3 g/L KH_2_PO_4_, and 18 g/L glucose adjusted to a pH of 7.2 with KOH) were inoculated with the overnight culture to an initial OD_600_ of 0.06-0.08 in Tunair shake flasks. Cells were grown at 37 °C and 250 rpm to an OD_600_ of 0.6 and inoculated with 0.5 mM isopropyl-β-D-thiogalactopyranoside (IPTG) to induce T7 RNA polymerase expression. At an OD_600_ of 3.0, cells were harvested by centrifugation at 5,000*g* for 10 minutes at 4°C. The resulting cell pellets were then resuspended with S30 buffer (10 mM Tris acetate pH 8.2, 14 mM magnesium acetate, and 60 mM potassium acetate) and pelleted by centrifugation at 10,000*g* for 2 minutes. Cells were washed a total of three times. After the final centrifugation step, the cell pellet mass was recorded and the cells were flash frozen in liquid nitrogen and stored at -80 °C.

Frozen cells were thawed on ice for 60 minutes and resuspended in 1 mL/g S30 buffer. Cells were then lysed with a single pass at 20,000-25,000 psi through either an Avestin EmusliFlex B15 or C3 homogenizer. The resulting lysate was centrifuged at 12,000*g* for 10 minutes and the supernatant collected. This step was performed twice. The final clarified lysate was aliquoted, flash frozen in liquid nitrogen, and stored at -80 °C.

#### Lysate Source Strain Engineering

The BL21 Star (DE3) Δ*gor* strain was constructed using the pcrEG and pEcCpfIH plasmids as previously described^100^. Initially, pEcCpfIH was introduced into BL21 Star (DE3) by chemical transformation. The strain containing the editing plasmid was utilized to prepare electro- competent cells and was induced with 50 mM arabinose at OD_600_ 0.1 and harvested at OD_600_ 0.6. Cells were washed with two washes of water and two washes of 10% w/v glycerol.

Concurrently, golden gate assembly was used to construct the cRNA expression plasmid, using the CRISPOR^101^ designed guide ATAGGAAGTATGAATACGGTCGA, targeting the *gor* gene. Homology directed repair (HDR) templates were designed containing 45 bp up- and downstream of the *gor* gene and ordered with phosphorothioate modified ends. Both the pcrEG- cRNA plasmid and the HDR templates were transformed into the prepared BL21 Star (DE3) strain containing the editing plasmid and recovered for 2-3 h at 37 °C before selection on LB agar containing 50 μg mL^-1^ kanamycin and 100 μg mL^-1^ spectinomycin.

Once colonies were confirmed for whole gene removal by colony PCR using Q5 hot start polymerase (NEB) and the primers gor-up-F 5’-ATT GAACTGGCGGTACTGCC-3’ and gor- down-R 5’-GTCAGAAGTACGGGTGGTGC-3’, the pcrEG-cRNA plasmid was removed by growing in 10 mM rhamnose overnight and streaked onto LB (no antibiotic) plates. Subsequently, the pEcCpfIH plasmid was removed by growing the strains overnight in LB containing 5 g L^-1^ glucose and streaked onto LB agar containing 5 g L^-1^ glucose and 10 g L^-1^ sucrose.

Bacterial genome sequencing was performed by Plasmidsaurus using Oxford Nanopore Technology with custom analysis and annotation.

#### DNA Template Preparation

All protein sequences used in this paper are listed in **Table S4**. Signal sequences and propeptides were removed when present to leave only the final, activated sequence. Unless otherwise noted, gene sequences were codon optimized for expression in *E. coli* and synthesized into a pJL1 backbone at the NdeI/SalI restriction sites by Twist Biosciences. Plasmids were purified for use in cell-free expression reactions with the Qiagen HiSpeed Plasmid Midi Kit and further cleaned with an ethanol precipitation.

#### Protein Purification

A plasmid containing DsbC with a 5’ CAT-Strep-linker (CSL) tag was transformed into NEB BL21(DE3) Competent *E. coli* cells and plated on LB agar containing 50 μg/mL kanamycin. The following day, a 50 mL culture of Overnight Express^TM^ Instant TB Media was inoculated with a single colony of transformed BL21(DE3) and grown overnight at 37 °C and 250 rpm. Cells were pelleted at 4,000*g* and 4 °C for 20 min. and resuspended in 2 mL of BugBuster® Master Mix. After incubation at room temperature for 15 min., the lysed cells were centrifuged at 10,000*g* for 10 min. to remove insoluble components. Meanwhile, 1 mL of Strep-Tactin®XT 4Flow® high-capacity resin was loaded onto a polypropylene column (Bio-Rad) and equilibrated with 2 column volumes of Buffer W (IBA). The lysed cell supernatant was added to the column and washed five times with 1 column volume of Buffer W. Protein was eluted with Buffer BXT (IBA). The most concentrated elution fractions were pooled and dialyzed into a buffer containing 100 mM Tris-Cl and 150 mM NaCl at pH 8. Protein yield and purity were assessed with a Bradford assay and SDS-PAGE gel. The purified protein was flash frozen in liquid nitrogen and stored at -80 °C.

#### Cell-free Gene Expression Reactions

10-15 μL cell-free reactions were performed at 30 °C in 2-mL microcentrifuge tubes (Axygen) for 20 h. All reactions contained 13.3 ng/μL plasmid and 30% v/v crude cell extract, unless otherwise noted. Reagent mixture compositions are detailed in **Table 1**. Reagents were thawed on ice and combined at room temperature to prevent precipitation of NMPs, which were always added to the reagent mixture last. To create an oxidizing reaction environment, cell extracts were treated with 500 μM or 25 μM iodoacetamide (IAM) at room temperature for 30 min. before use. An additional 4 mM oxidized glutathione (GSSG) and 1 mM reduced glutathione (GSH) were added to the reaction mixture. In some noted cases, 5 μM purified DsbC was also added to the reaction mixture^75,102^.

#### Bioreactor Tests

Custom, lab-made bioreactors (Sundberg *et al. In preparation*.) were rinsed with 70% ethanol, followed by deionized water, air-dried, assembled, and autoclaved with a 45-minute hold at 121 °C. After autoclaving, the reactors were cooled to 4°C for approximately 20 min. and then warmed to 30 °C in the reactor incubator for approximately 30 min. The components of the cell-free reagent mixture were thawed on ice, combined within a biosafety cabinet, and thoroughly vortexed between the addition of individual reagents. The reagent mixture and crude cell lysate were measured and aliquoted from a master mix container into 1.5-mL tubes for each of the three reactors and kept on ice until use. Dissolved oxygen and sfGFP fluorescence sensors were calibrated, and the reactors were flushed with the specified gas for approximately 10 min. prior to initiating the reactions. Cell-free reactions were initiated by drawing 2,800 μL of the reagent mix into a 5 mL syringe equipped with a blunt-tip 18-gauge needle and subsequently introduced into the reactor via a feed line. The impeller and sensors were activated as 1,200 μL of crude cell lysate was introduced into the reactor using a 3 mL syringe. The reactions were conducted for 20 h with a humidified gas flow of 400 cubic centimeters per minute into the reactor jacket, maintaining a constant impeller speed of 500 rpm and a temperature of 30 °C. After 20 h, the final cell-free reaction was harvested through the feed line, and the final reaction volume and sfGFP concentrations were assessed using a fluorescein standard.

#### Protein Expression Quantification

To assess the amount of sfGFP production, 2 μL of each cell-free expression (CFE) reaction were diluted with 48 μL of nanopure water in a black Corning Costar 96-well flat-bottom plate. Fluorescence was read with 485 nm excitation and 528 nm emission, with values converted to sfGFP concentration via a standard curve derived from sfGFP measured using ^14^C-leucine incorporation.

All other proteins were quantified using radioactivity, based on previously developed methods^103^. Briefly, 10 μM ^14^C-leucine was included in standard CFE reactions. After incubation, 5 μL of the total CFE reaction was treated with 100 μL 0.1 N KOH and incubated at 37 °C for 20 minutes. The remaining CFE reaction was centrifuged at 16,100*g* for 10 minutes, and 5 μL of the soluble supernatant was treated with KOH as well. 50 μL aliquots of the treated reactions were spotted onto two strips of Whatman 3MM CHR cellulose chromatography paper and dried under a heat lamp. One of the two chromatography paper strips was then placed in a beaker and washed three times with 5% w/v trichloroacetic acid for 15 minutes at 4 °C, followed by a wash with 200 proof ethanol at room temperature. The washed paper strips were dried under a heat lamp. Radioactivity was then measured with a Perkin Elmer MicroBeta2 with CytoScint liquid scintillation cocktail.

Autoradiograms were developed by separating total fractions of CFE reactions containing 10 μM ^14^C-leucine via SDS-PAGE. To visualize disulfide bond formation, samples were not denatured before analysis. Otherwise, samples were treated with dithiothreitol and denatured at 70 °C for 3 min. SDS-PAGE gels were vacuum-dried between two cellophane sheets with a Hoefer slab gel dryer and exposed to a phosphor screen for at least three days to generate autoradiographs. Autoradiogram gels were imaged with a Typhoon 7000. Protein bands were analyzed with densitometry.

Total yield of monoclonal antibodies was calculated by first calculating total yield from the ^14^C- leucine incorporation data assuming 100% heavy chain and 100% light chain production. These values were then averaged, due to the similar ratio of molecular weight to number of leucines. Full-length antibody yields were then calculated by multiplying this total yield by the percent full-length antibody calculated via densitometry from the oxidizing SDS-PAGE gels. Error was propagated through each step and visualized in plots with the error bars.

#### Protein Activity Assays

vtPA activity was determined by diluting cell-free reactions containing expressed vtPA 1:10 in PBS and incubating with 25.5 μM D-Ile-Pro-Arg-AMC peptide (iPR-AMC, Echelon Biosciences 855-18). Peptide cleavage by vtPA was determined by release of the free fluorescent 7-amino- 4-methylcoumarin (AMC) over the course of an hour at 26 °C and measured via a BioTek Neo2 plate reader with 354 nm excitation and 442 nm emission.

Colicin M antimicrobial activity was determined through incubation with CLM24 indicator cells, as previously described^50^. An overnight culture of LB media inoculated with CLM24 cells was diluted 1:100 in fresh LB media. The CLM24 culture was incubated at 37°C and 220 rpm to an OD_600_ of 0.7-0.9. The cells were then pelleted at 3,000*g* for 5 min., washed twice with 0.85% w/v sodium chloride, and resuspended in LB media to an OD_600_ of 0.1. Cell-free reactions containing diluted colicin M were added to 1.2 mL CLM24 cultures, incubated for 1 h at 37 °C and 220 rpm, and then washed twice with 0.85% w/v sodium chloride. The washed cell cultures were serially diluted, plated on LB agar, and incubated overnight at 30 °C before counting colonies.

TRI2-2 binding to the SARS-CoV-2 spike RBD protein was determined using AlphaLISA, based on previous descriptions^24^. AlphaLISA reactions were performed in a buffer containing 50 mM HEPES, 150 mM NaCl, 0.015% v/v TritonX-100, and 1 g/L BSA at pH 7.4. 2-μL reactions were prepared using an Echo 525 liquid handler, transferring solutions from an Echo Qualified 384- well PP PLUS plate (LabCyte PPL-0200) to a ProxiPlate 384-shallow well Plus plate (Revvity 6008280) using the 384_PP_Plus_AQ_GPSA fluid type. Serial dilutions of the His-tagged SARS-CoV-2 Spike RBD Protein (Acro Biosystems SPD-C82E9) and cell-free reactions incubated with either water or the TRI2-2_2xStrep plasmid were prepared using the AlphaLISA buffer. These dilutions were incubated with a final concentration of 0.02 mg/mL Anti-6xHis AlphaLISA acceptor beads (Revvity AL178C) for 1 h at room temperature before addition of Strep-Tactin AlphaLISA donor beads (Revvity AS106D) to a final concentration of 0.08 mg/mL. Following a second 1-h incubation, reaction luminescence was determined with a Synergy Neo2 plate reader with 80-ms excitation, 120-ms delay, and 160-ms integration time. Reactions were allowed to incubate for 10 minutes inside the plate reader before luminescence was recorded.

Trastuzumab binding was determined using a Trastuzumab Pharmacokinetic ELISA Kit (GenScript L00970). Cell-free reactions containing expressed trastuzumab were diluted 1:3,333 in PBS and assayed according to kit instructions.

#### Metabolite Analysis

Cell-free gene expression reactions were run to monitor changes in key metabolites over the course of the reaction. At each time point, a set of reactions in triplicate was quenched 1:1 with 10% trichloroacetic acid and flash frozen in liquid nitrogen. Samples were then thawed and centrifuged at 20,000*g* for 10 min. to remove precipitated protein. The supernatant was collected and 5 μL was injected onto an Agilent 1260 HPLC system. Metabolites were separated with an Aminex HPX-87H organic acids column at 60 °C with an isocratic flow of 5 mM sulfuric acid at 0.6 mL/min. Metabolite concentration was determined via refractive index detector based on the retention time and intensity of each compound’s standard solution. ATP was measured using the Promega CellTiter-Glo 2.0 Cell Viability Assay (G9241). Briefly, the quenched cell-free reaction supernatant was diluted 1:500 in nuclease-free water and mixed 1:1 with the CellTiter-Glo reagent; the luminescence was read on a BioTek Synergy H1 plate reader. ATP concentrations were determined based on a standard curve prepared with pure ATP. Inorganic phosphate concentrations were determined with the Sigma-Aldrich Phosphate Assay Kit (MAK308) using a 1:500 dilution of the quenched cell-free reaction supernatant.

Reaction pH was measured at the 15-μL scale using an Orion ROSS PerpHecT pH electrode. Cell-free reactions used for pH measurements were not quenched with trichloroacetic acid nor flash frozen in liquid nitrogen.

## Author Contributions

MLO, CEC, JRS, ASK, and MCJ contributed to the conceptualization of the study. MLO completed the majority of the research. RA prepared the genetically engineered lysate source strain. CAS performed the bioreactor experiments. ZMS assisted with the AlphaLISA experiments. MCJ, ASK, JRS, and GR supervised the research. MLO, ASK, and MCJ wrote the manuscript. All authors commented on and edited the manuscript.

## Data and materials availability

Source data for all figures are provided in the supplement.

